# Cell-type-resolved spatial proteogenomics from matched genome and proteome of the same cells

**DOI:** 10.64898/2026.07.09.737478

**Authors:** Maximilian Zwiebel, Maria Wahle, Rudolf Stadler, Mitchell P. Levesque, Reinhard Dummer, Thierry M. Nordmann, Matthias Mann

## Abstract

The genome and proteome of the same cells are rarely measured together, which is especially consequential in cancer, where somatic mutations vary across clones and drive disease. We show that a single standard proteomics extraction tip can retain peptides on-tip after digestion while genomic DNA passes into the normally discarded flowthrough. Combined with Deep Visual Proteomics, flowthrough co-isolation enables cell-type-resolved spatial proteogenomics from archival FFPE tissue, demonstrated in melanoma.

## Main text

Cells in a tissue share a genome yet differ vastly in the proteins they express. In cancer, somatic mutations accumulate into genetically distinct subclones whose early drivers and later alterations shape proteomic state, progression and therapy resistance. Spatially resolved methods such as 10x Visium^1^, MERFISH^2^, slide-DNA-seq^3^ or CODEX^4^ each capture only a single molecular layer. While technologies like 10x Chromium^5^ enable the parallel measurement of RNA and a panel of protein markers, the genome and proteome of the same cells have remained out of reach. Deep Visual Proteomics (DVP) resolves cell-type-specific proteomes in situ by AI-guided imaging and laser microdissection^6^, but not their matched genome. We reasoned that the matched genome could be recovered from a normally discarded fraction of the proteomics workflow itself.

In our standard MS-based proteomics workflow, we load a trypsin-digested lysate onto C18-based solid-phase extraction tips (Evotips) that retain peptides, while the peptide-depleted flowthrough is discarded^7^. We found that this flowthrough contains the sample’s genomic DNA in near-quantitative yield, so a single tip yields both peptides and DNA – an approach we term flowthrough co-isolation. Loading HeLa or melanoma cell digests onto Evotips and collecting the flowthrough recovered 85–95% of input DNA across roughly 50 to 1,000 cell equivalents (Fig. 1a,b; Methods). Fragments up to ∼10 kb passed cleanly during loading, while the subsequent wash contained only negligible amounts of DNA – a near-complete, single-pass separation (Fig. 1c). Sequencing libraries built from the flowthrough were indistinguishable from those built from unfractionated control DNA on a tapestation. To confirm that the flowthrough isolation does not introduce systematic bias in variant calling, we compared variant allele frequencies (VAFs) detected in matched A375 cell DNA and Evotip flowthrough, finding excellent concordance across heterozygous and homozygous variants (r = 0.99, J = 0.98; Fig. 1d). The bound peptide fraction was unaffected; the same tips delivered deep, low-input proteomes as expected of the DVP workflow, so neither measurement is traded against the other.

**Figure 1.**
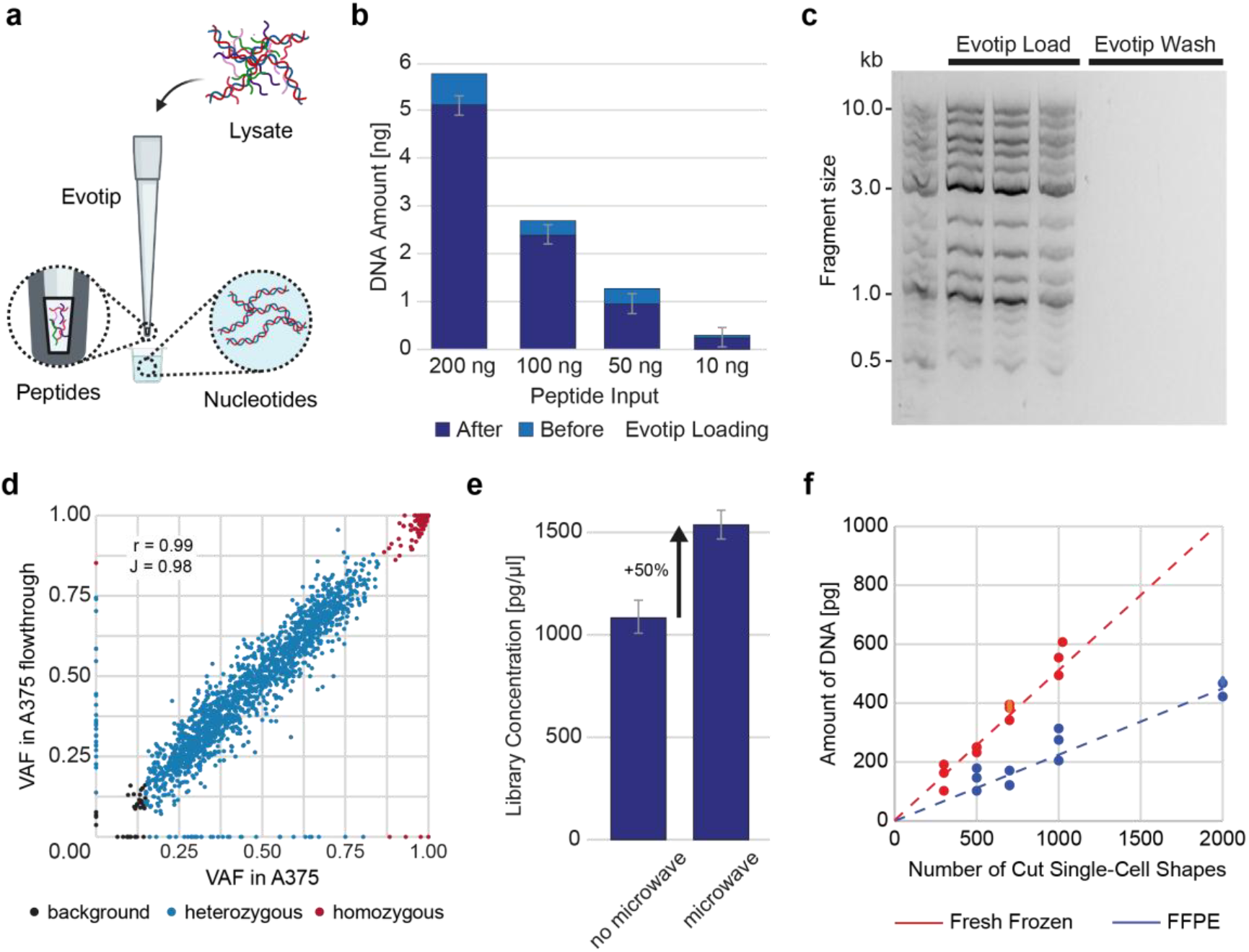
Flowthrough co-isolation of genome and proteome. (a) Schematic: peptides bind the Evotip C18 bed while genomic DNA passes into the flowthrough. (b) DNA measured before and after Evotip loading across a range of peptide inputs. (c) Agarose gel of load and wash fractions from an Evotip loaded with a DNA size ladder. (d) Variant allele frequencies (VAFs) from matched A375 genomic DNA versus Evotip flowthrough (r = 0.99, J = 0.98). Variants are color-coded by zygosity. (e) Library yield from FFPE flowthrough with and without microwave crosslink reversal. (f) DNA yield as a function of the number of laser-microdissected shapes for fresh-frozen and FFPE tissue.

Most clinical material is archived as formalin-fixed, paraffin-embedded (FFPE) tissue. While preserving the tissue architecture for decades^8^, the fixation procedure fragments nucleic acids and crosslinks them to proteins, rendering genomics from low-input FFPE samples challenging. We introduced a brief microwaving step to reverse formalin cross-links prior to flowthrough collection^9^, increasing library yield by roughly 50% (Fig. 1e). The tryptic digest performed routinely for proteomics further facilitates crosslink reversal, analogous to the proteinase K step used in dedicated FFPE DNA extraction protocols. Approximately 700 microdissected FFPE shapes (5 µm thickness) yielded ∼200 pg of DNA (equivalent to ∼300 fresh-frozen shapes), sufficient for whole-genome library preparation (Fig. 1f) – while the same material still delivered a deep proteome. The method is therefore applicable across input types, from bulk lysate to laser-microdissected tissue regions of only a few hundred cells, requiring no changes to the proteomics workflow apart from collecting the flowthrough.

Beyond providing a parallel genome, the mutation information from matched DNA should improve the proteome measurement itself. From the flowthrough sequencing we computed copy-number, single-nucleotide and structural variants using an adapted version of the MoCaSeq pipeline^10^ and built sample-specific protein sequence databases encoding each sample’s mutations and fusions (Fig. 2a; Methods). Searching against these personalized databases in DIA-NN^11^ identified 218 variant-only protein groups and 365 groups containing both canonical and variant proteoforms, sequences absent from a canonical database (Fig. 2b). Among these, we quantified variant peptides spanning mutated residues with high reproducibility across replicates (Fig. 2c), and demonstrated direct detection of mutation-bearing proteoforms at the peptide level (Fig. 2d). Because both derive from the same cells, genotype and its protein-level consequence are linked in one measurement, not inferred separately.

**Figure 2.**
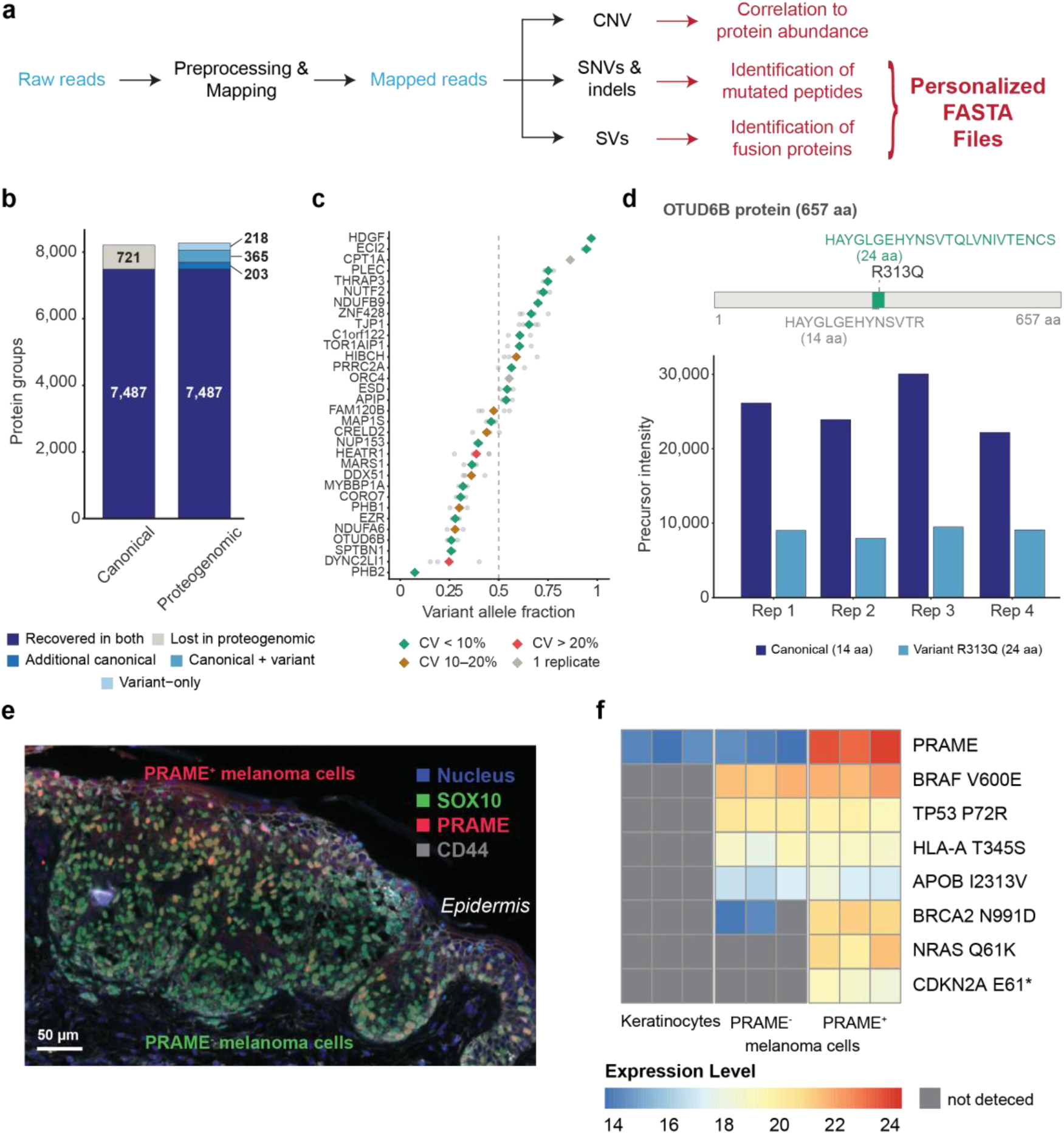
Cell-type-resolved spatial proteogenomics in melanoma. (a) Personalized database construction from variant and fusion protein sequences. (b) Protein-group identifications with personalized versus canonical databases (canonical, variant-only, and canonical + variant groups). (c) Variant allele fractions of peptides encoding SNVs from A375 cells, colored by coefficient of variation across replicates. (d) Variant versus canonical precursor intensities for the OTUD6B R313Q proteoform across four replicates. (e) Immunofluorescence of a melanoma section stained for SOX10, PRAME and CD44; PRAME-positive and PRAME-negative melanoma cells outlined for microdissection. (f) Proteomic levels of the detected variants across keratinocytes (control), PRAME-negative and PRAME-positive melanoma cells: BRAF V600E, NRAS Q61K and CDKN2A E61*.

We next applied flowthrough co-isolation, together with DVP, to human melanoma tissue, resolving matched genome and proteome within spatially defined cell populations in situ. PRAME-positive and PRAME-negative Sox10/CD44-positive melanoma cells were distinguished by a CD44/SOX10/PRAME imaging panel and AI classification before microdissection and captured separately (Fig. 2e; Methods). From these defined populations we detected the BRAF V600E driver in both, consistent with an early shared initiating event^12^, and identified NRAS Q61K and a truncating CDKN2A E61* alteration exclusively in the PRAME-positive subpopulation, matching the established temporal hierarchy of melanoma driver mutations^13^ – now resolved at the level of individual cell populations rather than bulk tissue (Fig. 2f). Beyond driver events, most detected variants represented passenger mutations or germline polymorphisms, typical of the mutational landscape of this tumor type and confirming that co-isolation captures the full spectrum of genomic variation without systematic bias.

By turning a discarded flowthrough into a matched genome, flowthrough co-isolation makes spatial proteogenomics broadly accessible, without specialized instrumentation or division of scarce material. Because it is independent of the capture chemistry and the downstream mass analyzer, the approach should extend to other extraction formats, to additional readouts such as RNA, and to the growing range of spatial and single-cell proteomic methods. With cohort sizes above a thousand^14^, the matched genotype-proteotype readout would enable intra-tissue pQTL analyses, linking somatic mutations to their protein-level consequences and dissecting the proteomic effects of cancer evolution at spatial resolution.

## Acknowledgements

This work was supported by the Max Planck Society for the Advancement of Science. We thank our colleagues at the Department of Proteomics and Signal Transduction at the Max Planck Institute of Biochemistry. In particular, Igor Paron and Tim Heymann for technical support as well as Medini Steger for administrative support. A special thanks goes to Rin Ho Kim from the Sequencing Core Facility at the Max Planck Institute of Biochemistry, who provided support for library sequencing.

## Methods

### Ethics and samples

Primary melanoma tissue biopsies were obtained during routine histopathological diagnostic procedures at the University Hospital Zurich. Retrospective analysis was performed with informed consent and ethical approval in place (Zurich: BASEC-Nr. 2022-01986). All experiments were performed in accordance with the Declaration of Helsinki. Histopathological diagnosis was re-validated by a board-certified dermatopathologist using H&E-stained tissue sections.

HeLa and A375 cells were cultured in Dulbecco’s Modified Eagle’s Medium (DMEM) supplemented with 10% fetal bovine serum (FBS) and 1% penicillin/streptomycin at 37°C in a humidified atmosphere with 5% CO_2_. Cells were cultured at 70–80% confluency and harvested by trypsinization. Cell pellets of 100,000 cells were collected by centrifugation and stored at −80°C until further processing.

### Membrane slide tissue mounting and immunofluorescent staining

PEN membrane slides (2 µm; MicroDissect GmbH) were UV-treated (254 nm, 1 h) and coated with Vectabond (Vector Laboratories, SP-1800-7) prior to tissue mounting. FFPE tissue blocks were sectioned at 5 µm using a microtome, mounted on pretreated slides, and dried overnight at 37°C. Sections were deparaffinized through sequential immersion in xylene (2 × 2 min) followed by graded ethanol washes (100%, 90%, 75%, 30%; 1 min each) and two rinses in distilled water. Antigen retrieval was performed in DAKO antigen retrieval solution (pH 9, S2367) supplemented with 10% glycerol at 92°C for 25 min in a water bath, followed by 20 min cooling at room temperature and two washes in distilled water. Sections were blocked with 5% BSA in PBS for 30 min at room temperature. Primary antibodies were applied as a cocktail in Dako Renoir Red diluent containing mouse anti-CD44 monoclonal IgG2a (abcam, clone 156-3C11, 1:200), mouse anti-SOX10 monoclonal IgG1 (NordicBiosite, bsh-7959-1, 1:200), and rabbit anti-PRAME monoclonal antibody (abcam, clone EPR20330, 1:100). CD44 was used as a cell surface and segmentation marker and SOX10 as a melanocytic lineage marker. Slides were incubated for 90 min at room temperature in a humidified chamber, washed twice in PBS (5 min each), and incubated with species- and isotype-specific secondary antibodies conjugated to Alexa Fluor dyes (anti-mouse IgG2a 568, anti-mouse IgG1 647, anti-rabbit 555; all Thermo Fisher, 1:400) for 60 min at room temperature. Nuclear counterstaining was performed with SYTOX Green (Thermo Fisher, 1:700 in distilled water, 7 min). After washing, slides were air-dried in darkness for 20–30 min and coverslipped with SlowFade Diamond antifade mountant (Thermo Fisher).

### Imaging, AI classification and laser microdissection

Stained slides were scanned on a Zeiss Axioscan 7 at 20× magnification with 10% tile overlap. Single-cell segmentation was performed in the Biological Image Analysis Software (BIAS) using a custom-trained Cellpose model. Only contours between 30 and 200 µm^2^ were retained. After removal of duplicates at tile-overlapping regions, a supervised machine learning classifier was applied to exclude false identifications and overlapping cell types. Three calibration points at characteristic tissue landmarks were exported together with contour coordinates. Contour outlines were simplified by removing 99% of data points prior to import into the LMD7 laser microdissection system (Leica) for semi-automated cutting with a 63× objective at the following settings: power 57, aperture 1–2, speed 23, middle pulse count 1, final pulse −3, head current 46–53%, pulse frequency 2,600, offset 210. PRAME-positive and PRAME-negative tumor-cell populations, as well as keratinocytes, defined as CD44 positive and SOX10/PRAME-negative cells within the epidermis, were collected into separate wells of a 384-well plate (Eppendorf).

### Proteomics sample preparation

For cell line samples, cell pellets of 100,000 cells were resuspended in 100 µl lysis buffer (10% acetonitrile, 60 mM TEAB in mass spectrometry-grade H2O) and lysed at 72°C for 30 min in a thermal shaker (Eppendorf, 1,200 rpm). Samples were digested overnight with 200 ng trypsin and 400 ng LysC at 37°C and quenched the following day with 10 µl of 10% formic acid (approximately 1% final concentration). Peptide concentrations were determined by tryptophan fluorescence assay prior to loading.

For microdissected DVP shapes, collection wells were washed with 28 µl of 100% acetonitrile and dried in a SpeedVac for 20 min at 45°C. Samples were resuspended in 5 µl lysis buffer and sealed with adhesive foil. For FFPE-derived material, a microwave cross-link reversal step was applied (2 min at 400 W, 2 min at 800 W). Samples were then heated at 72°C for 90 min in a 384-well thermal cycler (Eppendorf) for cell lysis. Overnight digestion was performed with 4 ng trypsin and 6 ng LysC at 37°C and quenched after 18 h with 10 µl buffer A (0.1% formic acid).

### Flowthrough co-isolation of DNA and peptides

Digested cell and tissue lysates were loaded onto Evotip Pure tips (Evosep). Tips were first soaked in 1-propanol for 3 min and primed twice with 50 µl of buffer B (acetonitrile with 0.1% formic acid), followed by a second 1-propanol soaking step for 1 min. Tips were then equilibrated twice with 50 µl of buffer A (0.1% formic acid). Samples were loaded and the flowthrough was captured in a 96-well plate (Eppendorf) for subsequent library preparation. Tips were washed once with buffer A and stored with 150 µl buffer A on top until LC-MS measurement. All centrifugation steps were performed at 700 × g for 1 min.

To characterize the DNA fragment size permeability of the Evotip C18 bed, tips were prepared as described above, loaded with 5 µl of a DNA size standard (1 kb Plus DNA Ladder, NEB), and loading and wash fractions were collected separately. Fractions were analyzed by gel electrophoresis on a 1.5% agarose gel (120 V, constant voltage).

### Mass spectrometric acquisition

Peptides were separated on an Evosep One LC system (Evosep Biosystems) using the Whisper 40 samples per day (SPD) method with an Aurora Elite CSI third-generation C18 column (15 cm × 75 µm ID, 1.7 µm; IonOpticks) at 50°C inside a nanoelectrospray ion source (Captive Spray, Bruker Daltonics). Mobile phases were 0.1% formic acid in LC-MS-grade water (buffer A) and acetonitrile / 0.1% formic acid (buffer B). The timsTOF SCP (Bruker Daltonics) was operated in diaPASEF mode with an optimal acquisition method generated using py_diAID, consisting of 8 diaPASEF scans with variable window widths and 2 ion mobility windows per scan, covering an m/z range of 300–1,200 and an ion mobility range of 0.7–1.3 Vs cm^−2^. The instrument was operated in high sensitivity mode with accumulation and ramp times of 100 ms, a capillary voltage of 1,400 V, and a collision energy applied as a linear ramp from 20 eV at 1/K0 = 0.6 Vs cm^−2^ to 59 eV at 1/K0 = 1.6 Vs cm^−2^.

### DNA library preparation and sequencing

Prior to library preparation, flowthroughs were purified and buffer-exchanged using AMPure XP beads (Beckman Coulter; 0.8x bead ratio, elution in 13 µl 10 mM TE buffer). Purified fractions were used for sequencing library preparation using the NEBNext Ultra II FS kit (NEB). DNA was fragmented and end-repaired by incubation with the NEBNext Ultra II FS Enzyme Mix in the supplied reaction buffer (37°C for 5 min, followed by 65°C for 30 min). Sequencing adapters (NEBNext, diluted to 0.6 µM in 10 mM TE buffer) were ligated using the NEBNext Ultra II Ligation Master Mix with Ligation Enhancer (20°C, 15 min), followed by USER enzyme treatment to cleave adapter hairpins (37°C, 15 min). Size selection and cleanup were performed using AMPure XP beads with a double size-selection protocol (0.4x followed by 0.2x bead ratio). Libraries were amplified by PCR using NEBNext Ultra II Q5 Master Mix and TruSeq primers (13 cycles). Final libraries were purified using AMPure XP beads (0.8x) and eluted in 30 µl Buffer EB. All working volumes were halved compared to the manufacturer’s manual.

Library size distributions were analyzed on a TapeStation (Agilent) and quantified using a Qubit fluorometer (Thermo Fisher). Samples were measured on an Illumina NovaSeq 6000 platform (Illumina) in 2x 150 bp paired-end mode to 20x coverage.

### Genomic data analysis

Genomic variants were called using an adapted version of the MoCaSeq pipeline^10^, with all steps run against the human reference genome (GRCh38.p14). In brief, raw reads were trimmed and aligned to the reference genome, followed by postprocessing and base quality score recalibration. Somatic SNVs and indels were called using Mutect2, either in matched tumor–normal mode for microdissected tissue samples with available matched normal, or in tumor-only mode with germline filtering for cell line samples. Copy-number variations were detected using HMMCopy and structural variants with Delly. Variants were annotated using VEP and SnpEff to obtain protein-level consequence predictions. A custom Python script was then used to parse the resulting MAF files, extract amino acid changes, and write mutant protein sequences to a DIA-NN-compatible FASTA file for use as sample-specific proteogenomic search databases.

### Proteomics data analysis

Raw data were analyzed using DIA-NN 2.5. The search was done without fixed modifications; variable methionine oxidation was omitted to maintain Proteoforms scoring mode and minimize search space inflation. For DVP tissue samples, per-sample FASTA files were generated by concatenating the canonical UniProt human reference proteome with sample-specific variant sequences derived from MoCaSeq variant calls and searched using Proteoforms scoring and --pg-level 1. Variant protein sequences were encoded as individual FASTA entries with UniProt-compatible headers carrying unique accessions encoding the amino acid substitution (e.g. sp|P15056_V600E|BRAF_HUMAN). For cell line samples, a two-search strategy was employed: a first search against the canonical UniProt proteome alone to generate a standard proteome quantification matrix, and a second search against the combined canonical and variant FASTA to enable variant peptide detection and allelic ratio quantification. Variant peptide identifications were extracted from the precursor-level report, retaining only proteotypic variant peptides detected in at least two replicates. Allelic ratios were computed by summing precursor intensities across charge states for each variant and its canonical counterpart, with variant allele fraction defined as variant intensity divided by the sum of variant and canonical intensities.

### Statistical data analysis

DNA recovery was calculated as the ratio of DNA amount measured in the flowthrough after Evotip loading to the total input DNA, quantified by fluorometric assay (Qubit, Thermo Fisher). All DNA quantification measurements are presented as mean with 95% confidence intervals across biological replicates. VAF concordance between matched A375 genomic DNA and Evotip flowthrough was assessed by Pearson correlation coefficient (r) and Jaccard similarity index (J). Variants were classified by zygosity using a maximum likelihood classifier based on the binomial distribution, evaluating each variant under three hypotheses, background noise, heterozygous, and homozygous alternate, given the observed read depth and number of alt-supporting reads. Variants with a posterior probability below 0.9 for the assigned class were flagged as ambiguous and excluded from down-stream analyses. Protein groups from canonical and proteogenomic DIA-NN searches were compared by matching leading protein accessions; groups were classified as recovered in both, lost in the proteogenomic search, additional canonical, canonical + variant, or variant-only based on their presence and accession composition across searches. Coefficient of variation (CV) for allelic ratios was calculated across four replicates. All data analysis and visualization was performed in R (v4.5.1).

